# A realistic in-silico brain phantom for quantifying susceptibility anisotropy-induced error in susceptibility separation

**DOI:** 10.64898/2026.04.07.716972

**Authors:** Daniel Ridani, Benjamin De Leener, Eva Alonso-Ortiz

## Abstract

**Purpose:** To create a realistic in-silico brain phantom for positive and negative magnetic susceptibility that incorporates susceptibility anisotropy, enabling the evaluation of how susceptibility anisotropy influences susceptibility separation algorithm performance.

**Methods:** We expanded an existing QSM validation phantom by creating separate maps for positive and negative susceptibility, with the option of modeling susceptibility anisotropy. Multi-echo gradient echo data were simulated to evaluate four susceptibility separation techniques (χ-separation, DECOMPOSE-QSM, APART-QSM, and 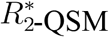). To assess the impact of noise, simulations were performed at different SNR levels (50, 100, 200, 300).

**Results:** Our findings showed that the error in negative susceptibility estimates increased by up to 53% when susceptibility anisotropy was present, compared to the case without susceptibility anisotropy, with χ-separation being the algorithm that was most sensitive to anisotropy. Robustness to noise varied across the assessed algorithms, with APART-QSM and χ-separation having the highest and lowest sensitivity to noise, respectively.

**Conclusion:** The modified phantom is open-source and can serve as a numerical ground truth for evaluating susceptibility separation methods. Our findings emphasize the importance of incorporating susceptibility anisotropy into susceptibility separation models to improve their accuracy.

## Introduction

Quantitative susceptibility mapping (QSM) is a quantitative MRI (qMRI) technique that measures the underlying magnetic susceptibility (χ) distribution of biological tissues^1^. QSM has emerged as a valuable technique for evaluating changes in iron and myelin within gray matter (GM) and white matter (WM)^2–4^. These changes are linked to natural aging and neurodegenerative diseases such as multiple sclerosis (MS), Parkinson’s disease (PD), and Alzheimer’s disease (AD). For example, in aging, a biphasic susceptibility pattern in white matter tracts (e.g., the internal capsule) illustrates how myelin initially develops (lowering susceptibility) and later degenerates (raising susceptibility)^3,5^. In neurodegenerative diseases, QSM has been increasingly used as a biomarker to track pathological changes. In MS, elevated susceptibility arises from both demyelination and iron deposition due to inflammation^6–8^. In PD, increased susceptibility in the substantia nigra correlates with dopaminergic cell loss and overall disease severity^9,10^. Similarly, QSM studies in early-stage AD have revealed susceptibility differences in deep gray matter and cortical regions compared to healthy controls^11,12^.

While QSM is commonly used to assess iron content, what is in fact measured by QSM is the bulk magnetic susceptibility arising from the sum of all sources within a voxel. A high positive measured χ value would typically be interpreted as being due to a high iron concentration. Yet, iron and myelin are frequently co-localized in the brain. This means that with QSM, decreases in myelin (which has negative, χ^−^, susceptibility) cannot be discerned from increases in iron (which has positive, χ^+^, susceptibility), and vice versa. Recently, efforts have been made to separate the contributions of positive and negative magnetic susceptibility sources to measured signals (here we refer to any such approach as “susceptibility separation”). Advances by Shin et al. (χ-separation)^13^, Chen et al. (DECOMPOSE-QSM)^14^, Li et al. (APART-QSM)^15^, and Dimov et al. 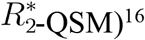 rely upon biophysical models to extract susceptibility source concentrations from MRI 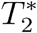 or *T*_2_ and 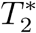 signals. In healthy volunteers^13^, χ-separation χ^+^and χ^−^ maps show distributions that track those of histologically confirmed iron and myelin distributions in the brain. In another recent application of χ-separation, Lee et al.^17^ showed that iron and myelin (inferred from χ^+^ and χ^−^ measurements) peak at different laminar surfaces of the cortex with different intensity profiles and that QSM laminar profiles are similar to those of χ^+^ measurements but with different peaking points, seemingly due to a bias from myelin. This work shows the exciting promise of susceptibility separation for studying the role of iron and myelin in aging and disease. However, one must consider that a basic assumption of both QSM and current susceptibility separation methods is that the macroscopic susceptibility in an imaging voxel is isotropic. This assumption holds true for GM^18^. Conversely, several studies have shown that the χ of WM depends on the orientation of axons with respect to the main magnetic (*B*_0_) field^19–21^, suggesting that χ of WM is in fact anisotropic. The susceptibility anisotropy of brain WM has been attributed to myelin, in particular to the highly ordered lipid molecules within the myelin sheath^4,22^.

QSM involves various complex processing steps^23^: (1) gradient-echo (GRE) phase unwrapping, (2) creation of a field map from the unwrapped phase, (3) removal of the background field to obtain a field map that reflects only local sources of susceptibility, and (4) dipole inversion to result in a local χ map. Numerous approaches have been proposed over the years for each one of these steps^24,25^ and significant strides have been made by the community to validate and standardize QSM. Historically, physical phantoms such as iron and calcium mixtures have been used to mimic positive and negative susceptibility sources. While these phantoms are useful for reproducing bulk susceptibility contrasts, they remain limited by their isotropic magnetic behavior and cannot represent susceptibility anisotropy. Digital phantoms overcome many of the constraints of physical phantoms by providing anatomical realism with accurate tissue boundaries, partial volume effects, background field structure and the possibility to model susceptibility anisotropy. They also allow controlled variation of imaging parameters such as B_0_ strength, SNR, echo time, repetition time, and flip angle, which would be time consuming to vary and rescan using a physical phantom. In addition, they can be easily shared, making them reproducible and easily extendable. The latter supports consistent benchmarking and the future exploration of additional features, thereby providing a comprehensive platform for evaluating QSM acquisition and processing strategies. Given these advantages, the Electro-Magnetic Tissue Properties Study Group (EMTP-SG) of the International Society of Magnetic Resonance in Medicine (ISMRM) recommends that the assessment of QSM processing methods be conducted using a digital phantom (to serve as a numerical ground truth)^26^. In support of this, a digital QSM validation head phantom^27^ was recently developed and used to establish consensus guidelines for the acquisition and processing of QSM data. The QSM validation phantom includes 7 T quantitative *T*_1_ and 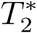 maps, M_0_, and χ, as well as diffusion tensor imaging (DTI) data that can be used to simulate MR signals for testing of QSM acquisition protocols and data processing methods. The phantom cannot, however, be used to test susceptibility separation techniques.

To address this limitation, here we propose to extend the QSM validation phantom so that it can be used to validate susceptibility separation algorithms. Our proposed phantom can be used for 3 T and 7 T MRI simulations and includes isotropic χ^+^ and isotropic or anisotropic χ^−^ values. We demonstrate the utility of our proposed phantom by using it to simulate realistic MRI data, which is subsequently used to compare the results of four^13–16^ susceptibility separation algorithms: χ-separation, DECOMPOSE-QSM, APART-QSM, and 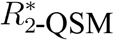. Our comparisons provide insights into the impact of susceptibility anisotropy on separation algorithm performance.

## Methods

### Phantom creation

The original QSM validation phantom was amended to replace voxelwise χ with χ^+^ and χ^−^ values. Given that χ-separation and APART-QSM require *T*_2_ maps (which are not provided by the QSM validation phantom), 3 T and 7 T *T*_2_ maps were created. Additionally, the original 7 T QSM validation phantom *T*_1_ map was converted to 3 T, enabling MR simulations at both field strengths.

#### Tissue segmentation

Data analyses were conducted using the regions of interest (ROIs) provided with the QSM validation phantom (Table 1, column 1) and additional white matter sub-ROIs that we introduced (Table 1, column 2). White matter sub-ROIs were manually segmented using ITK-Snap^28^, guided by a color-coded fractional anisotropy (FA) map.

**Table 1.** List of ROIs included in the QSM validation phantom (column 1) and newly added WM sub-ROIs (column 2).

| QSM validation phantom ROIs | Additional WM sub-ROIs |
| --- | --- |
| Caudate nucleus | Body of the corpus callosum |
| Globus pallidus | Splenium of the corpus callosum |
| Putamen | Genu of the corpus callosum |
| Red nucleus | Anterior limb of the internal capsule |
| Dentate nucleus | Posterior limb of the internal capsule |
| Substantia nigra | Superior corona radiata |
| Thalamus | Posterior corona radiata |
| White matter | Anterior corona radiata |
| Grey matter | Posterior thalamic radiations |
|  | Superior longitudinal fascicle |

#### Susceptibility maps

For each QSM validation phantom ROI, piecewise (i.e., uniform) χ^+^ and χ^−^ values were assigned to replace that of the bulk susceptibility (χ^*total*^) included in the QSM validation phantom. χ^+^ values were first assigned by ensuring consistency with χ^+^ values obtained from a χ-separation atlas^29^:

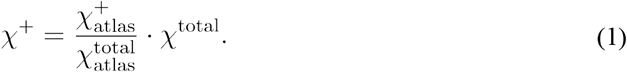

Once the χ^+^ values were calculated, their correlation with histology-derived iron concentrations reported in literature^30^ was computed (*r* = 0.83).

The negative susceptibility (χ^−^) was then calculated by subtracting the positive susceptibility from the total susceptibility: χ^−^ = χ^*total*^-χ^+^.

To incorporate realistic intensity modulations (i.e., texture) into the phantom, piecewise susceptibility values for each ROI (Table 1, column 1) were modulated according to^27^:

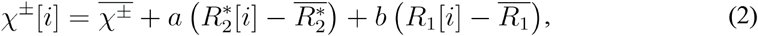

where *i* denotes the voxel index and 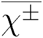 is the mean positive or negative susceptibility value across each ROI. The parameters *a* and *b* are proportionality constants that determine the weighting of the 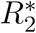 and *R*_1_ relaxation rate variations within each ROI. These parameters were set to the same values used in the QSM validation phantom^27^. The terms 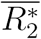 and 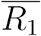 are the ROI-averaged apparent transverse and longitudinal relaxation rates, respectively. For the χ^−^ map, we set *b* = 0, so that only demeaned 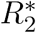 modulates texture; for the χ^+^ map, we set *a* = 0 to avoid introducing the orientation-dependent anisotropy of 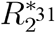 into χ^+^. We demeaned 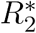 and *R*_1_ so that texture has zero mean within each ROI, thereby preserving ROI-average susceptibility while adding realistic spatial variation. This texturing step is included only to avoid piecewise-homogeneous ROIs and to generate more visually realistic maps. It is not derived from, nor intended to approximate, established susceptibility–relaxation theory and therefore should not be interpreted as implying proportional or biophysical relationships between susceptibility and relaxation maps.

Our phantom has the option to account for WM’s magnetic susceptibility anisotropy. For this, we replaced the isotropic χ^−^ with the apparent magnetic susceptibility^20^:

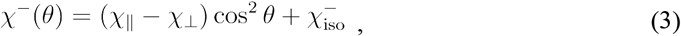

where χ∥ and χχ are tensor components representing the susceptibility of myelinated fibers parallel and perpendicular to their principal axes, respectively, *θ* is the fiber-to-field angle derived from the QSM validation phantom DTI data using the primary eigenvector map (V1)^32^, and 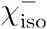 represents any orientation-independent susceptibility. In Sibgatulin *et al*.^*33,34*^, the authors measured the difference between χ∥ and χχ (denoted *δ*χ) and 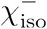 using single-orientation acquisitions. In Li *et al*.^*20*^, the authors measured χ∥ and χχ individually through multiple head orientation acquisitions. Using the average reported values by Sibgatulin *et al*. and Li *et al*., we assigned piecewise *δ*χ and 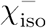 values to each WM sub-ROI. The generated *δ*χ and χ_iso_ maps were then weighted in each WM sub-ROI using the *R*_1_ map (which was additionally subjected to Gaussian noise (*η*) (mean = -0.04,

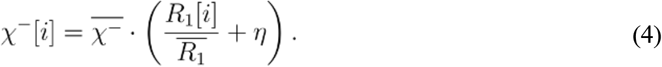

standard deviation = 0.05, to result in a more realistic intensity modulation): The final (ROI-averaged) susceptibility values are listed in Table 2.

**Table 2.**
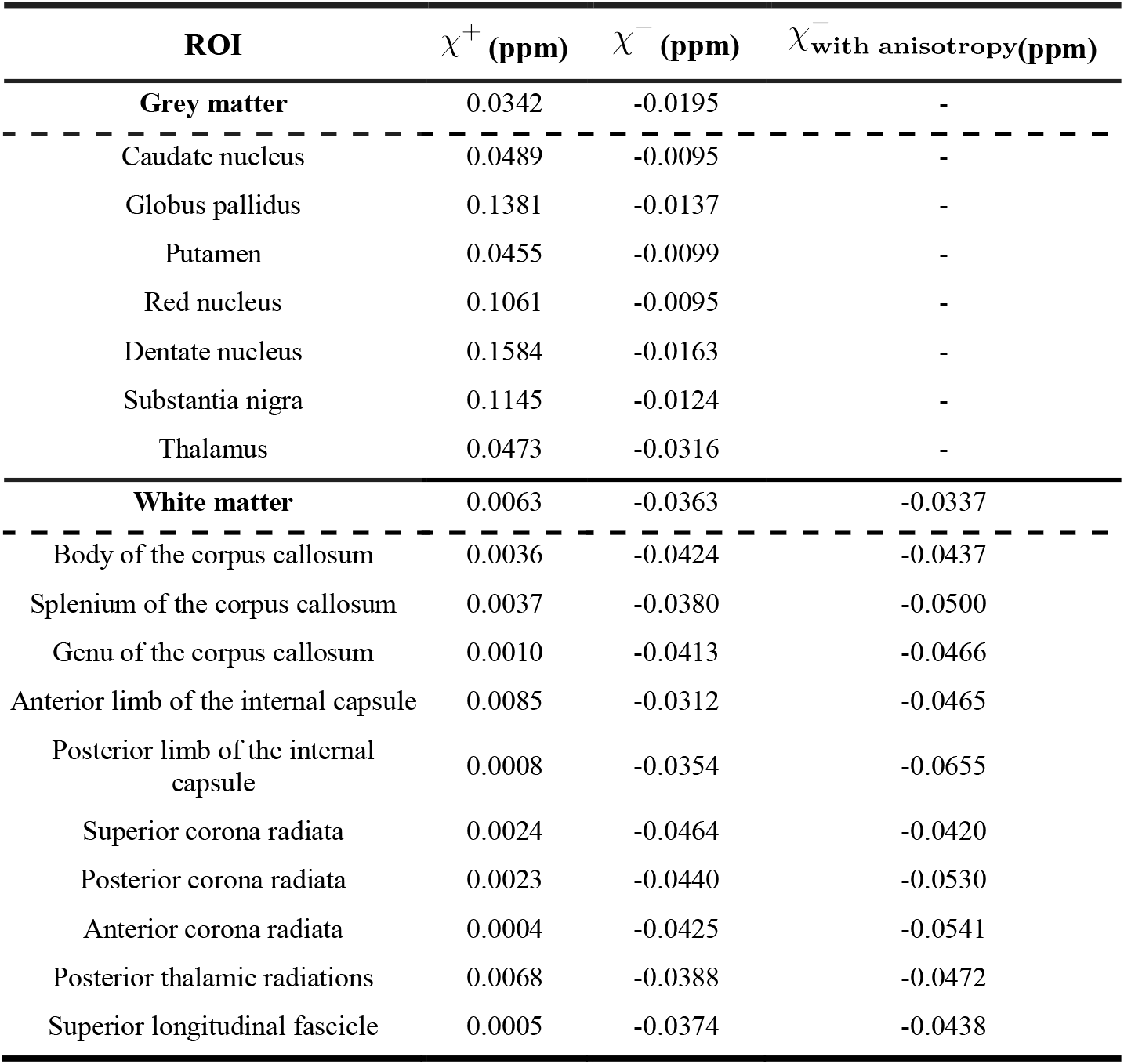
ROI-averaged magnetic susceptibility ( χ^+^, χ^−^, and 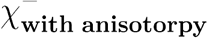) after applying a realistic intensity modulation.

#### Relaxation maps

A 3 T *T*_2_ map was created by assigning 3 T literature-reported *T*_2_ values to each QSM validation phantom ROI^35,36^. To transform these piecewise *T*_2_ values into a realistic map, we used the 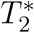 and *M*_0_ maps available in the QSM phantom dataset. For each ROI, we computed the mean 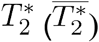 and the mean 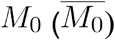. We then normalized each voxel’s 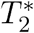 and *M*_0_ values within the ROI by dividing them by their regional means. The voxel-wise *T*_2_ values were then modulated using normalized relative offsets according to the equation:

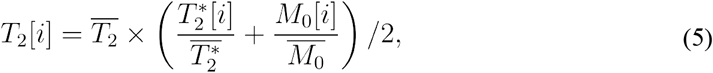

where 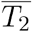 is the mean (piecewise) *T*_2_ value assigned to the region. It should be noted that the texturizing steps introduce realistic intra-regional variability while preserving ROI-mean values, and do not imply biological causation.

Additionally, a 7 T *T*_2_ map was generated by scaling the 3 T *T*_2_ map. Cox et al.^37^ reported *T*_2_ values for multiple ROIs at 3 T and 7 T. To derive the relationship between *T*_2_ values at these field strengths, we computed the ratio *T*_2_(7 T) / *T*_2_(3 T) for each ROI. The average of these ratios was then used to generate a scaling factor. The ratio showed limited variability across GM and WM (mean = 0.65, SD = 0.03). This approach was adopted instead of directly using the literature-reported 7 T *T*_2_ values due to the absence of corresponding measurements for all ROIs listed in Table 1, column 1. As a result, the following equation was derived to estimate *T*_2_ at 7 T based on *T*_2_ at 3 T:

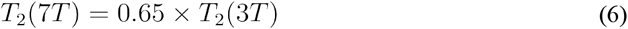

To estimate *T*_1_ values at 3 T, we employed a model described by Bottomley *et al*.^*38*^:

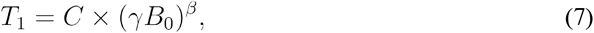

where *γ* is the gyromagnetic ratio and 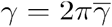, with 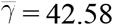 MHz/T. *C* and *β* are tissue-specific parameters. Rooney *et al*.^*39*^ provided *T*_1_ values for multiple regions across a range of magnetic field strengths from 0.15 T to 7 T and obtained the *C* and *β* estimates by fitting the model to (total) WM and GM regions. Using Rooney *et al*.’s reported *T*_1_ values for the ROIs included in our phantom (Table 1, column 1), non-linear least squares fitting was performed to determine each ROI’s optimal *C* and *β* values. *C* and *β* values are reported in the Supporting Information Table S1 for all Table 1, column 1 ROIs.

### Data simulation

Spoiled gradient-recalled-echo data was simulated using the steady-state equation:

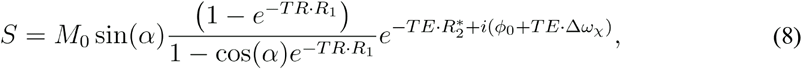

where *α* is the flip angle, TR and TE are the repetition and echo times, respectively, *ϕ*_0_(*r*) is the initial phase, and Δ*ω* _χ_is the frequency shift due to susceptibility differences.

Δ*ω* _χ_ is related to the magnetic field perturbation, Δ*B*, in the following way:

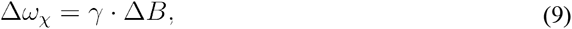

Δ*B* can be predicted by convolving the dipole kernel, *D*, with the local magnetic susceptibility offset, Δ _χ_ :

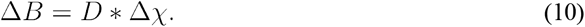

Incorporating positive and negative susceptibility effects into the GRE signal requires considering the relationship between 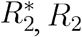, and 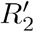. The total decay rate 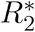 is given by:

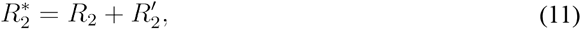

where 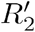 accounts for the susceptibility-related signal decay. 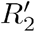 term is dependent on the total susceptibility χ^total^, which includes both χ^+^ and χ^−^ susceptibility components:

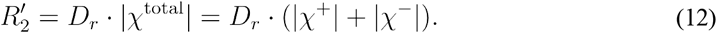

*D*_*r*_ is a susceptibility-dependent relaxation constant influenced by the shape of the susceptibility source^40,41^. Positive susceptibility sources, such as iron, are modeled as a sphere: 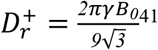, and negative susceptibility sources are modeled as parallel cylinders: 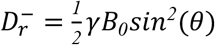, where *θ* is the angle between the cylinder and the main magnetic field^41^. The theoretical value for 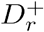 in GM is 107.83 Hz/ppm/T, while for the cylindrical model in WM, 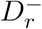 ranges from 0 to 133.76 Hz/ppm/T, for *θ* = 0 to *θ* = 90◦, respectively. Thus, the complete expression for 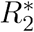 becomes:

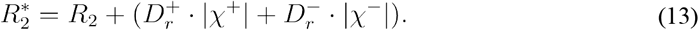

Combining Eq. 8, Eq. 9, Eq. 10, and Eq. 13, the final signal equation is:

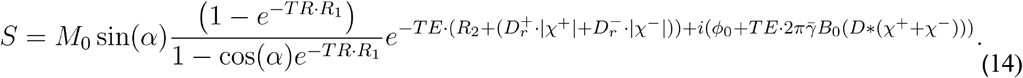

#### 3 T simulations

To evaluate the impact of susceptibility anisotropy on susceptibility separation results, we conducted noiseless simulations with and without susceptibility anisotropy effects. Using Eq. 14, our phantom’s χ^+^ and χ^−^ maps, and the *T*_1_, *T*_2_, and D_r_ maps specific to 3 T, we simulated 3 T MGRE magnitude and phase data with TR = 50 ms, flip angle = 15°, and TE_1_/ΔTE/TE_6_ = 3/4/23 ms. Furthermore, to assess the algorithm’s robustness to noise, we performed simulations (accounting for susceptibility anisotropy) at SNR = [50, 100, 200, 300] by adding Gaussian noise with a mean value of zero to the GRE signal’s real and imaginary components^42^.

#### 7 T simulations

Noiseless MGRE magnitude and phase were also simulated at 7 T with susceptibility anisotropy effects, using our phantom’s χ^+^ and χ^−^ maps, and the *T*_1_, *T*_2_, and D_r_ maps specific to 7 T. The imaging parameters were TR = 50 ms, flip angle = 15°, and TE_1_/ΔTE/TE_4_ = 4/8/28 ms. These parameters were suggested by the QSM validation phantom for 7 T simulations^27^.

### Susceptibility separation algorithm validation

Figure 1 illustrates the workflow for validating susceptibility separation algorithms. Simulated 3 T and 7 T MGRE magnitude data were first fit to a mono-exponential decay model, resulting in an 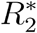 map. Then, an 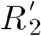 map was calculated by subtracting the *R*_2_ map from the 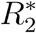 map. MGRE phase data underwent a series of processing steps to be usable for susceptibility separation algorithms: Laplacian phase unwrapping^43^ was applied, and the variable-kernel sophisticated harmonic artifact reduction for phase data (VSHARP) background field removal method^44^ with a spherical mean value of 12 was used to compute the local field map. Because one of the susceptibility separation algorithms (DECOMPOSE-QSM) required a QSM image as input, rather than a local field map, we used FANSI^45^ for dipole inversion to generate the necessary QSM image.

**Figure 1.**
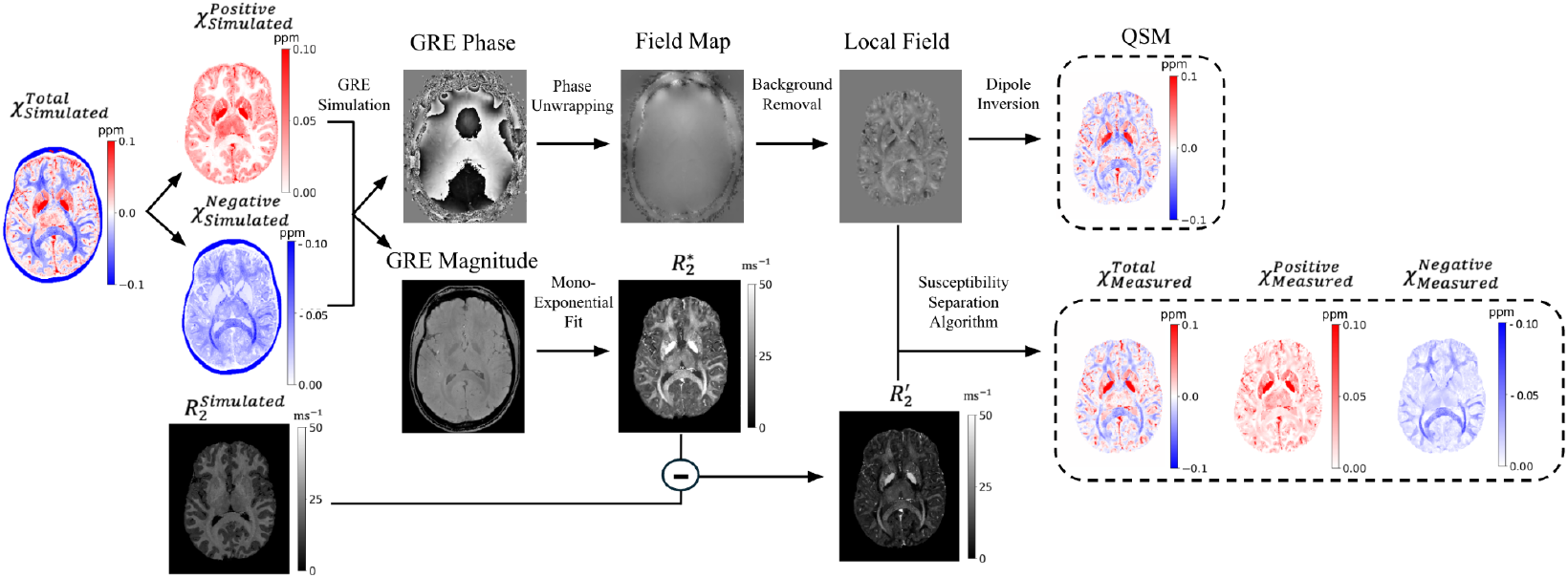
Workflow schematic. Magnitude GRE data was used to compute 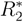 through a mono-exponential fit. Phase GRE data underwent a series of processing steps (phase unwrapping, background field removal, and dipole inversion) to generate local field and QSM maps. In addition, the reversible transverse relaxation rate 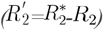 was computed. Finally, the local field and 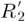 maps were input into susceptibility separation algorithms.

The processed 3 T and 7 T data were fit to four open-access susceptibility separation algorithms: χ-separation^13^, 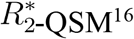, APART-QSM^15^, and DECOMPOSE-QSM^14^, resulting in χ^+^ and χ^−^ maps.

First, we assessed whether there were systematic differences between the 3 T and 7 T fit results for χ^−^ with susceptibility anisotropy, using a Bland-Altman analysis of values averaged within WM sub-ROIs (Table 1, column 2). Following this, all subsequent investigations were conducted exclusively at 3 T. To examine the impact of susceptibility anisotropy, susceptibility separation χ^−^ results obtained from noiseless data both with and without the inclusion of susceptibility anisotropy effects were averaged within WM sub-ROIs. For each sub-ROI, the mean squared percentage error (MSPE) between measured and simulated χ^−^ values was computed.

Next, to investigate the sensitivity to noise of the different susceptibility separation algorithms, we computed the average MSPE between simulated and measured χ^+^ and χ^−^ for each ROI in Table 1 with susceptibility anisotropy effects and varying degrees of SNR. To avoid bias due to differences in ROI sizes, the MSPE was normalized by the volume of the corresponding ROI before averaging. The ROI-normalized MSPE values were then averaged across ROIs.

To further evaluate the influence of noise in the presence of susceptibility anisotropy effects, additional WM ROIs were created. These WM ROIs were obtained by first co-registering the anatomical image of a WM fiber-bundle atlas^46^ to our simulated MGRE magnitude TE1 image using an affine transformation. The resulting transformation was subsequently applied to the atlas’s WM fiber-bundle segmentation labels. Manual corrections were then performed to refine the co-registered segmentation labels. The latter was used to mask our phantom’s fiber orientation (*θ*) map, resulting in a *θ* map that only contains fiber angles for voxels that correspond to major fiber-bundles. Finally, voxels were binned into 10-degree intervals, resulting in new WM ROIs that are categorized according to their average *θ*. The use of the fiber-bundle segmentation labels was intended to ensure that ROIs with *θ*≅0 originate from fibers that are truly parallel to *B*_0_, as opposed to WM regions with many crossing fibers. Next, for each WM bundle bin, the MSPE between the simulated and measured χ^−^ values was calculated. To ensure that extreme voxel values did not bias these regional error estimates, outliers were removed using an interquartile range (IQR)-based filter (1.5×IQR criterion) applied within each bin prior to averaging. This process was repeated across all noise levels (SNR 50, 100, 200, and 300). Finally, the maximum error variation (MEV) was calculated to quantify the difference in error for ROIs with fibers oriented parallel (MSPE_0_o) and perpendicular (MSPE_90_o) to *B*_0_ . The MEV quantifies the extent to which noise affects the variability in error between these two extreme orientations:

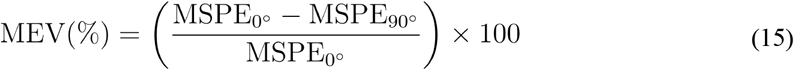

### Susceptibility separation phantom validation using in-vivo data

We acquired in-vivo data from a healthy volunteer (age = 22 years, female) on a 3 T Prisma Siemens MRI scanner. By comparing these in-vivo data with our phantom’s data, we sought to confirm that the phantom effectively mimics real tissue behavior. The volunteer provided informed consent, and the study was approved by the Comité d’éthique de la recherche du Regroupement Neuroimagerie Québec.

A 3D multi-echo gradient-echo (MGRE) sequence was acquired with the following parameters: field of view (FOV) = 192 × 192 × 134 mm^3^, voxel size = 0.7 × 0.7 × 0.7 mm^3^, number of slices = 160, repetition time (TR) = 27 ms, echo times (TE) = 3 ms, 7 ms, 11 ms, 15 ms, 19 ms, and 23 ms, bandwidth = 420 Hz/pixel, flip angle = 12°, phase partial Fourier = 6/8, GRAPPA = 2, and total acquisition time = 8.5 minutes. For *T*_2_ mapping, a 2D accelerated turbo spin-echo (TSE) sequence^47^ was used with the following parameters: FOV = 192 × 192 × 111 mm^3^, voxel size = 0.8 × 0.8 × 1.5 mm^3^, number of slices = 74, TR = 4860 ms, echo times from TE_1_ = 10 ms to TE_6_ = 60 ms with an echo spacing (ΔTE) of 10 ms, bandwidth = 977 Hz/pixel, flip angle = 180, GRAPPA= 2, and total acquisition time = 10.5 minutes.

For tissue segmentation, a T1-weighted anatomical image was acquired using a MP-RAGE sequence with the following acquisition parameters: FOV = 224 × 224 × 264 mm^3^, voxel size = 1.0 × 1.0 × 1.0 mm^3^, number of slices = 176, TR = 2400 ms, TE = 2.17 ms, inversion time TI = 1000 ms, bandwidth = 210 Hz/pixel, flip angle = 8°, GRAPPA = 2, total acquisition time = 5.5 minutes.

The MGRE data were processed by applying Laplacian phase unwrapping^43^ to the phase images, followed by VSHARP^44^ background field removal. The FANSI^45^ dipole inversion algorithm was used to compute QSM maps, used as an input for DECOMPOSE-QSM. Magnitude images from the MGRE data were fit to a mono-exponential decay model to derive 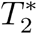 maps. For the multi-echo spin-echo (MESE) data, mono-exponential fitting was performed to obtain *T*_2_ maps, which were subsequently used to calculate 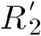 maps. The processed data were then fit to χ-separation, 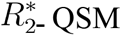, APART-QSM, and DECOMPOSE-QSM.

FSL’s segmentation tool “FAST”^48^ was used to segment (with some manual corrections) the ROIs listed in Table 1 (column 1) from the T1-weighted image. Measured χ^+^ and χ^−^ values were averaged within the ROIs and used to compute the correlation with the susceptibility separation phantom’s matching ROI-averaged χ^+^ and χ^−^ values. Additionally, in-vivo and simulated local field maps were compared within the predefined WM sub-ROIs (Table 1). The correlation coefficient and slope of the linear regression were calculated to assess the phantom’s ability to reproduce the field variations characteristic of anisotropic tissues. Both in-vivo and simulated local field maps were referenced to whole-brain average field offsets. Finally, the agreement between 3 T in-vivo and generated *T*_2_ maps was assessed within the same ROIs described above. For visual comparisons with in-vivo data, our simulated MGRE magnitude TE1 was co-registered to the in-vivo MGRE magnitude TE1 using an affine transformation. The resulting transformation was then applied to the phantom’s susceptibility separation results, local field map, and *T*_2_ map, which were subsequently resampled to match the spatial resolution of the in-vivo data.

## Results

### Susceptibility phantoms and relaxation maps

Figure 2 shows the simulated χ^+^ and χ^−^ maps (under “Susceptibility Phantom”) both with and without the effects of susceptibility anisotropy. Directional variations in susceptibility, indicative of the anisotropy effect, are visible in the χ^−^ map when anisotropy effects are included. Under “Maps”, top row, is the generated 3 T *T*_2_ map and the 7 T *T*_2_ map extrapolated from 3 T. Below those images is the 3 T *T*_1_ map extrapolated from 7 T, and the 7 T *T*_1_ map originating from the QSM validation phantom. Under “ROI”, we show the QSM validation phantom ROIs and the WM sub-ROIs used for analyses.

**Figure 2.**
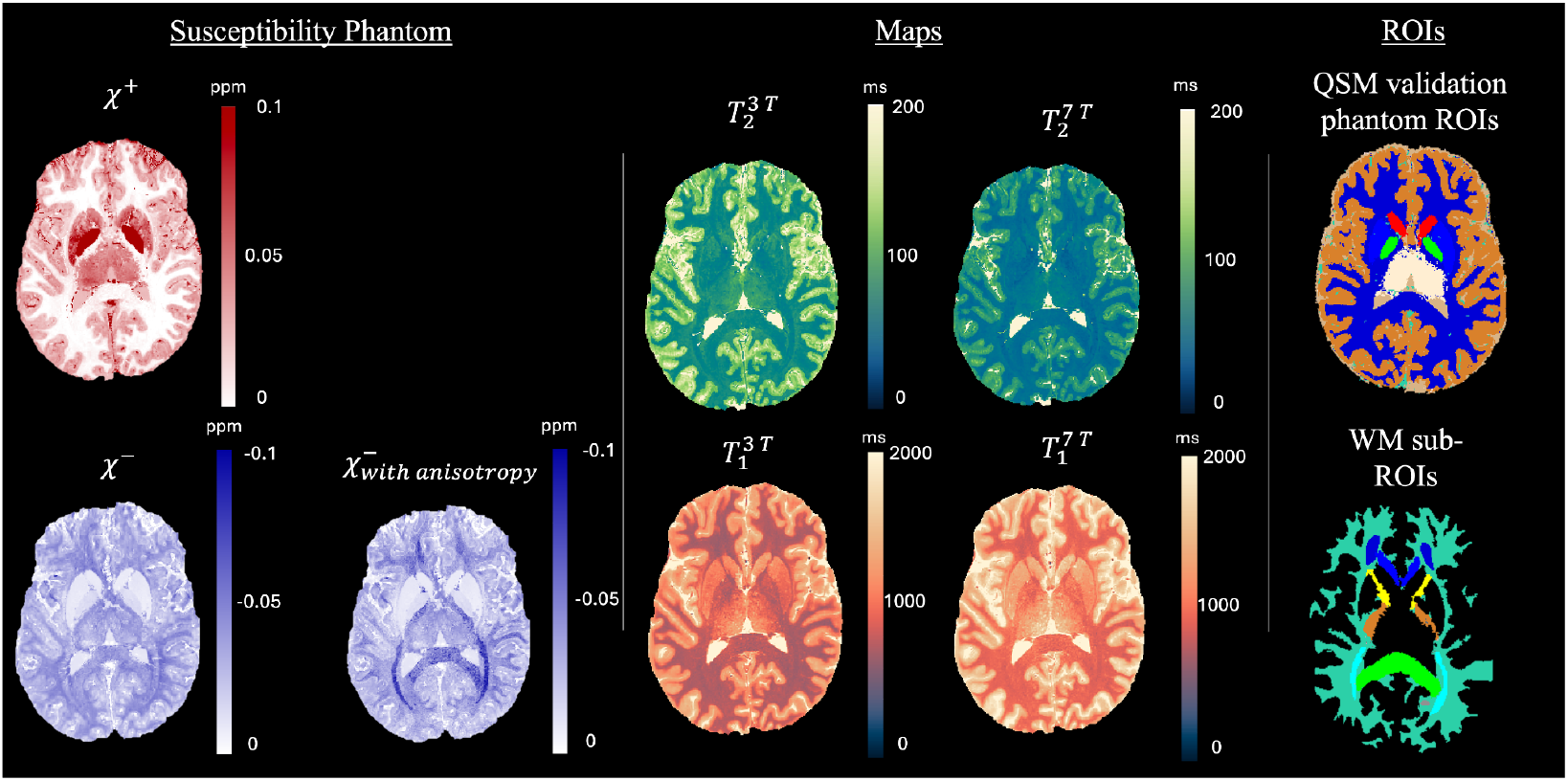
Left: susceptibility maps (χ^+^, χ^−^, and 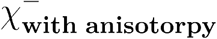); Middle: T_2_ and T_1_ Maps for 7 T and 3 T; Right: QSM validation phantom ROIs and WM sub-ROIs.

Table 2 presents the ROI-averaged χ^+^, χ^−^, and 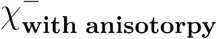 (after applying a realistic intensity modulation). χ^+^ values in the deep grey matter structures exhibit the highest susceptibility χ^−^ values, consistent with their high iron content. Conversely, χ^−^ values are the lowest in the WM ROIs, reflecting their high myelin content.

### Impact of susceptibility anisotropy

Bland-Altman analyses (Supplementary Material, Figure S1) indicated a weak systematic difference between 3 T and 7 T measured χ^−^ with susceptibility anisotropy effects included. Given these findings, we subsequently focused on 3 T simulations. Figure 3 shows kernel density estimate (KDE) plots of the WM sub-ROI-averaged MSPE values for χ^−^ measured from noiseless 3 T data simulated with and without susceptibility anisotropy effects. Without anisotropy, χ-separation demonstrates the lowest mean MSPE at 10.4%, compared to APART-QSM, 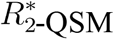, and DECOMPOSE-QSM, with MSPEs of 10.9%, 14.2%, and 15.0%, respectively. When anisotropy is introduced, APART-QSM scored the lowest MSPE at 12.9%, compared to χ-separation, 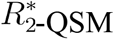, and DECOMPOSE-QSM, with MSPEs of 16.0%, 15.5%, and 17%, respectively. As seen in Figure 3, for every algorithm tested, the distribution of MSPE with anisotropy appears to be broader and has a higher average than that of the case without anisotropy. More specifically, the relative increase in MSPE due to the inclusion of anisotropy was 53% (unpaired t-test with a degree of freedom df = 18, t-value = 4.164 *p* = 0.0003) for χ-separation, 18% for APART-QSM (df = 18, t-value = 1.841, *p* = 0.075), 10% for 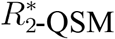 (df = 18, t-value = 0.473, *p* = 0.638), and 13% for DECOMPOSE-QSM (df = 18, t-value = 1.113, *p* = 0.272). The comparison of χ^+^ values with and without anisotropy is shown in the Supporting Information Figure S2 and S3.

**Figure 3.**
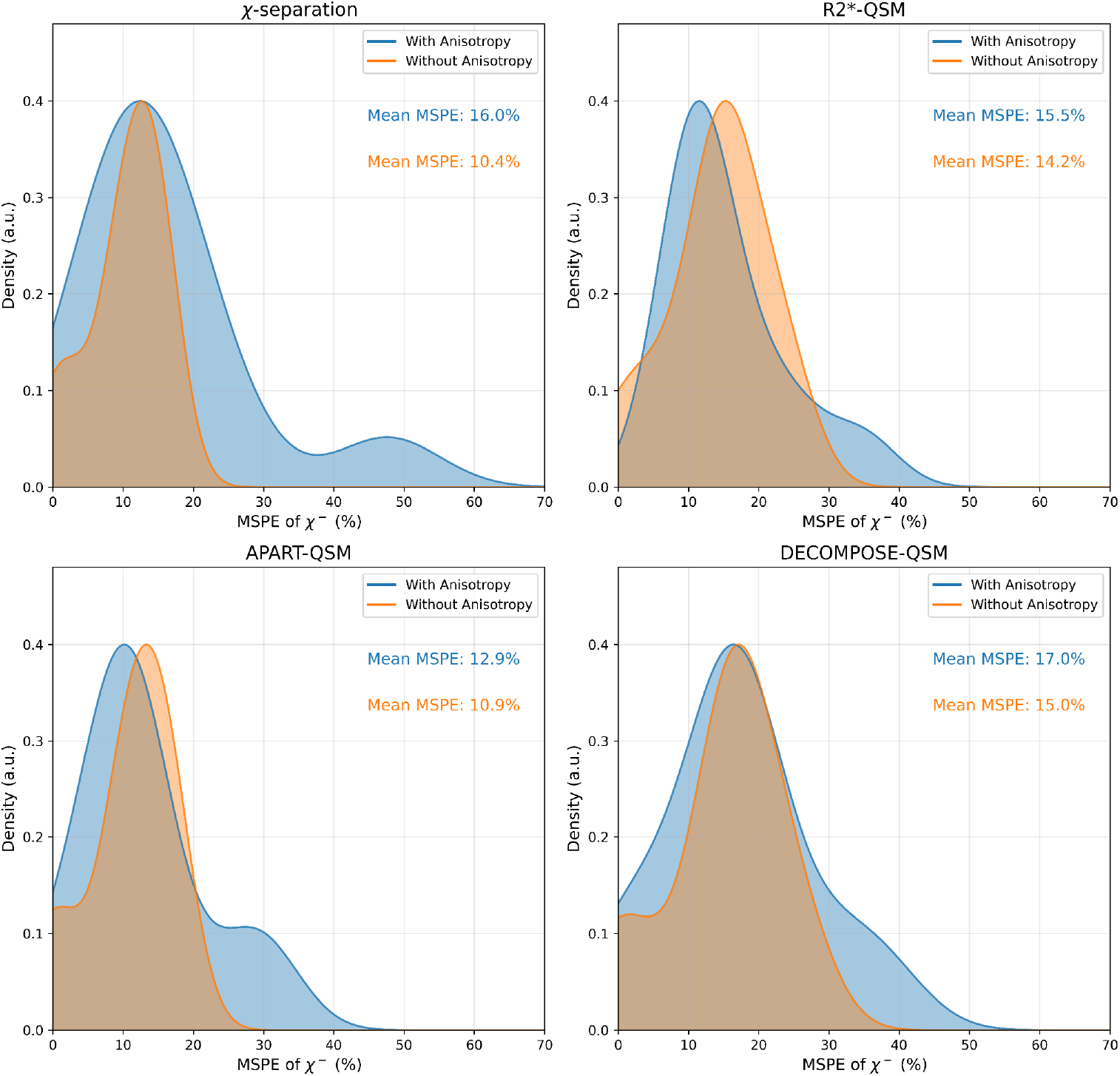
KDE plots of WM sub-ROI-averaged MSPE of χ^−^ measured from noiseless 3 T data simulated with (blue) and without (orange) susceptibility anisotropy using four susceptibility separation techniques: χ -separation, 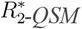, APART-QSM, and DECOMPOSE-QSM.

### Impact of GRE noise

Figure 4 illustrates the ROI-averaged MSPE of χ^+^ and χ^−^ obtained with four different susceptibility separation methods and GRE data simulated at SNR = [50, 100, 200, 300]. χ - separation consistently shows the lowest MSPE for χ^+^ estimates, decreasing from approximately 21% at SNR 50 to around 9% at SNR 300. 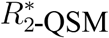 follows a similar trend, with MSPE values dropping from 33% to 19% as SNR increases, while APART-QSM sees the steepest decrease, with over 90% MSPE at SNR 50; but reaching a comparably low MSPE of around 14% at SNR 300. DECOMPOSE-QSM also follows a similar pattern, with MSPE decreasing from close to 74% at SNR 50 to around 21% at SNR 300.

**Figure 4.**
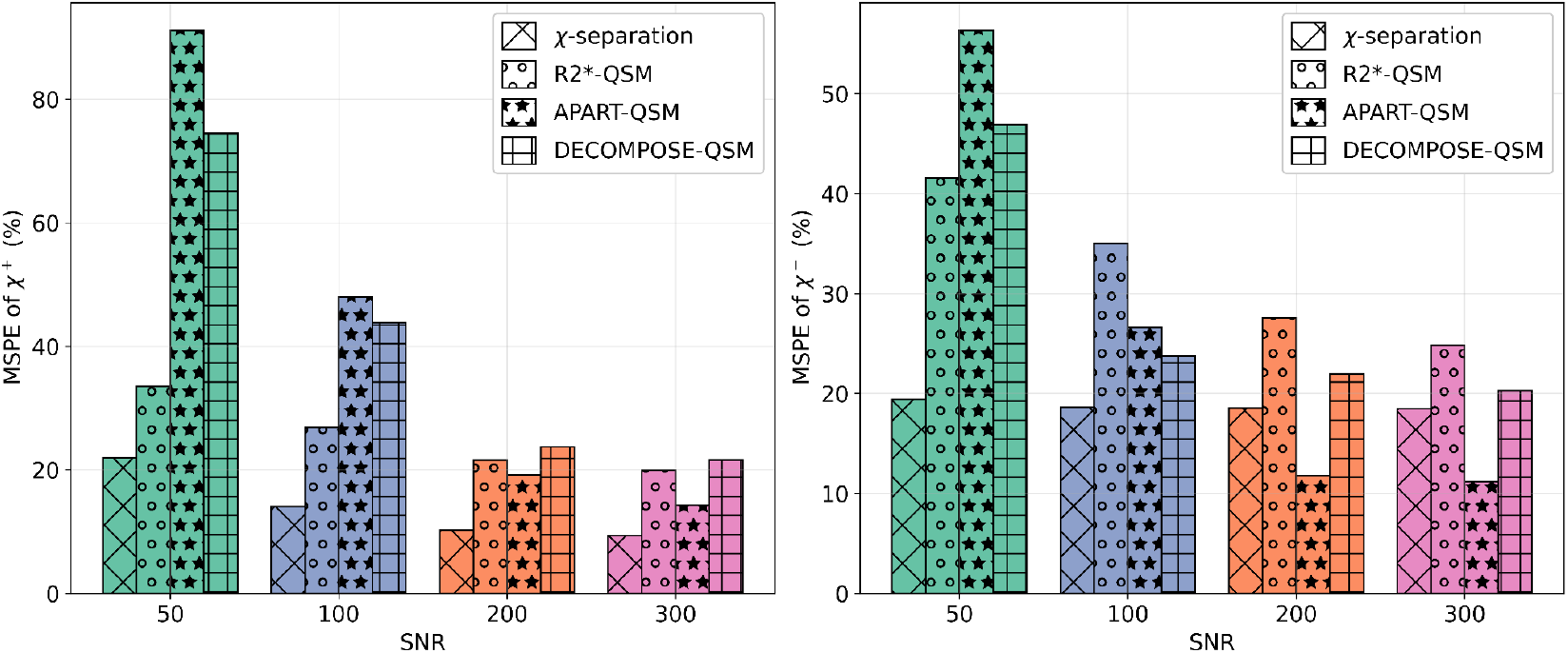
MSPE across varying signal-to-noise ratios (SNR) forχ -separation, 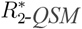, APART-QSM, and DECOMPOSE-QSM. The left panel displays the MSPE for χ^+^ and the right panel shows the MSPE for χ^−^ .

For the χ^−^ component, APART-QSM initially achieves the highest MSPE at SNR 56 (about 60%), but its error drops rapidly as SNR increases, reaching around 11% at SNR 300. χ-separation maintains stable performance, with MSPE gradually dropping from 19% to 17% as SNR increases. Both 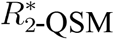 and DECOMPOSE-QSM display similar behaviors, starting with high MSPE at low SNR (MSPE around 41% and 46%, respectively), but experiencing a decrease at high SNRs, where 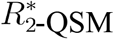 reaches nearly 25% and DECOMPOSE-QSM approaches 20%.

In Figure 5, the MSPE of χ^−^ was averaged across WM voxels partitioned into 10° *θ-*bins ranging from 0° to 90°, for the four different susceptibility separation methods and SNR level = [50, 100, 200, 300]. For all methods, MSPE is highest when *θ* is low, where the effect of anisotropy is maximized, and MSPE is lowest at high *θ*. For high SNR values (SNR = 300), the plots exhibit pronounced variation with orientation angle, reflecting the impact of anisotropy. As SNR decreases, the error percentage increases and the curves flatten, highlighting how the anisotropy effect becomes less noticeable in noisy conditions. The MEV values support this observation, with higher SNR values associated with larger variation. For χ-separation, the MEV decreases from 68.0% at SNR 300 to 26.3% at SNR 50. Similarly, for 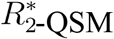, the MEV declines from 53.7% to 35.2%. For APART-QSM, the MEV drops from 50.0% at SNR 300 to 9.9% at SNR 50, while DECOMPOSE-QSM shows a reduction from 37.4% to 21.9%.

**Figure 5.**
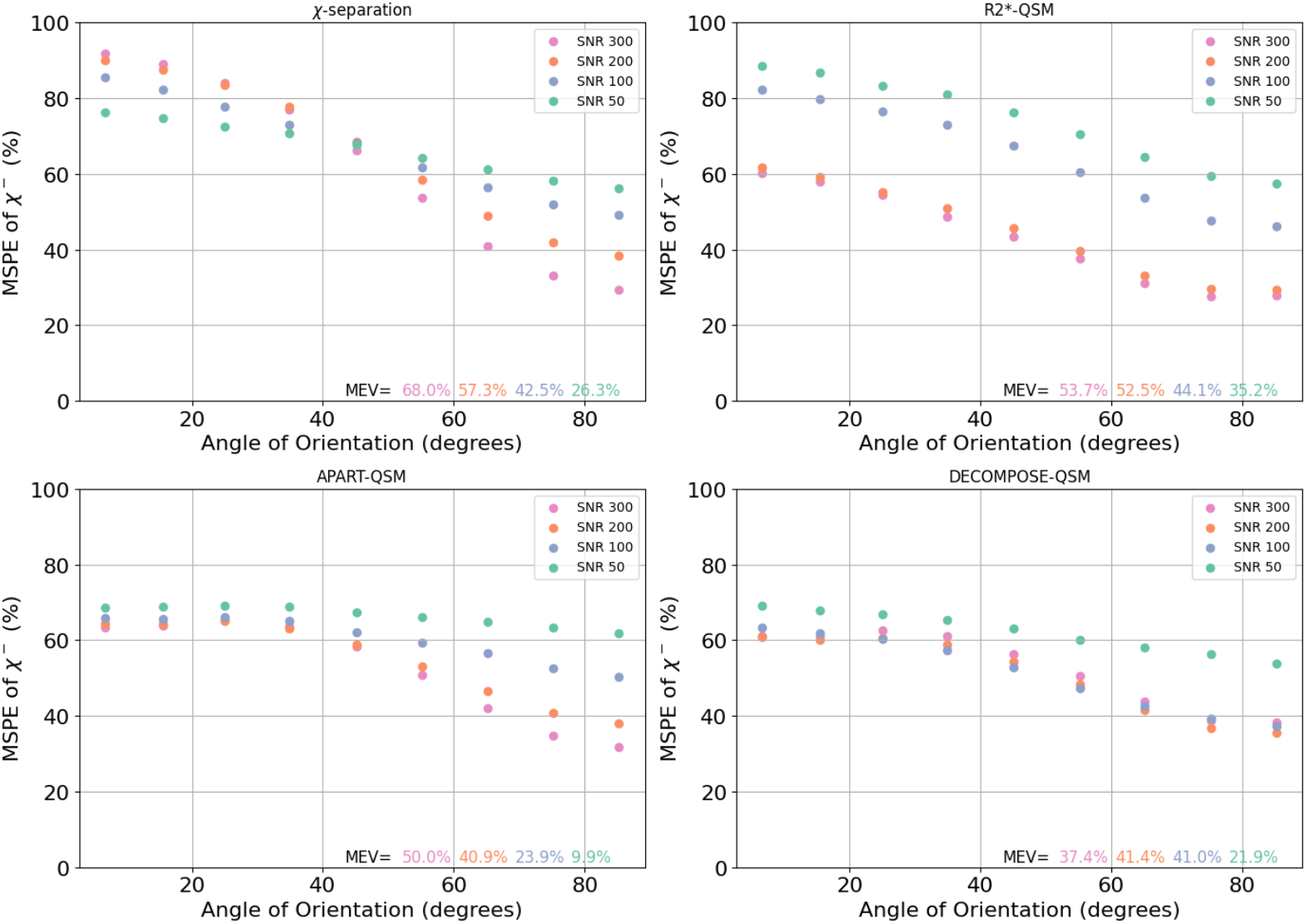
MSPE as a function of the WM sub-ROIs θ (in degrees) for different SNR levels (300, 200, 100, and 50) alongside the MEV for each SNR. The data points were obtained by calculating the MSPE for each angle of orientation, sampled at 10-degree intervals, for four algorithms: -separation (top-left), 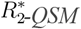 (top-right), APART-QSM (bottom-left), and DECOMPOSE-QSM (bottom-right).

### Comparison between simulated vs. in-vivo susceptibility maps

To confirm that the phantom accurately mimics in-vivo data, we compared simulated and in-vivo susceptibility values (shown in Figure 6). Qualitative comparisons of simulated and in-vivo measured χ^+^ and χ^−^ (Figure 6A) across the four susceptibility separation algorithms revealed similar contrast and spatial patterns, suggesting good agreement between the two. Quantitatively, strong correlations were observed between the simulated and in-vivo measured susceptibility values (Figure 6B). For χ^+^, all algorithms demonstrated very strong correlations (*r* > 0.94). Correlations for χ^−^ were slightly lower, but remained strong (*r* > 0.76) across all methods.

**Figure 6.**
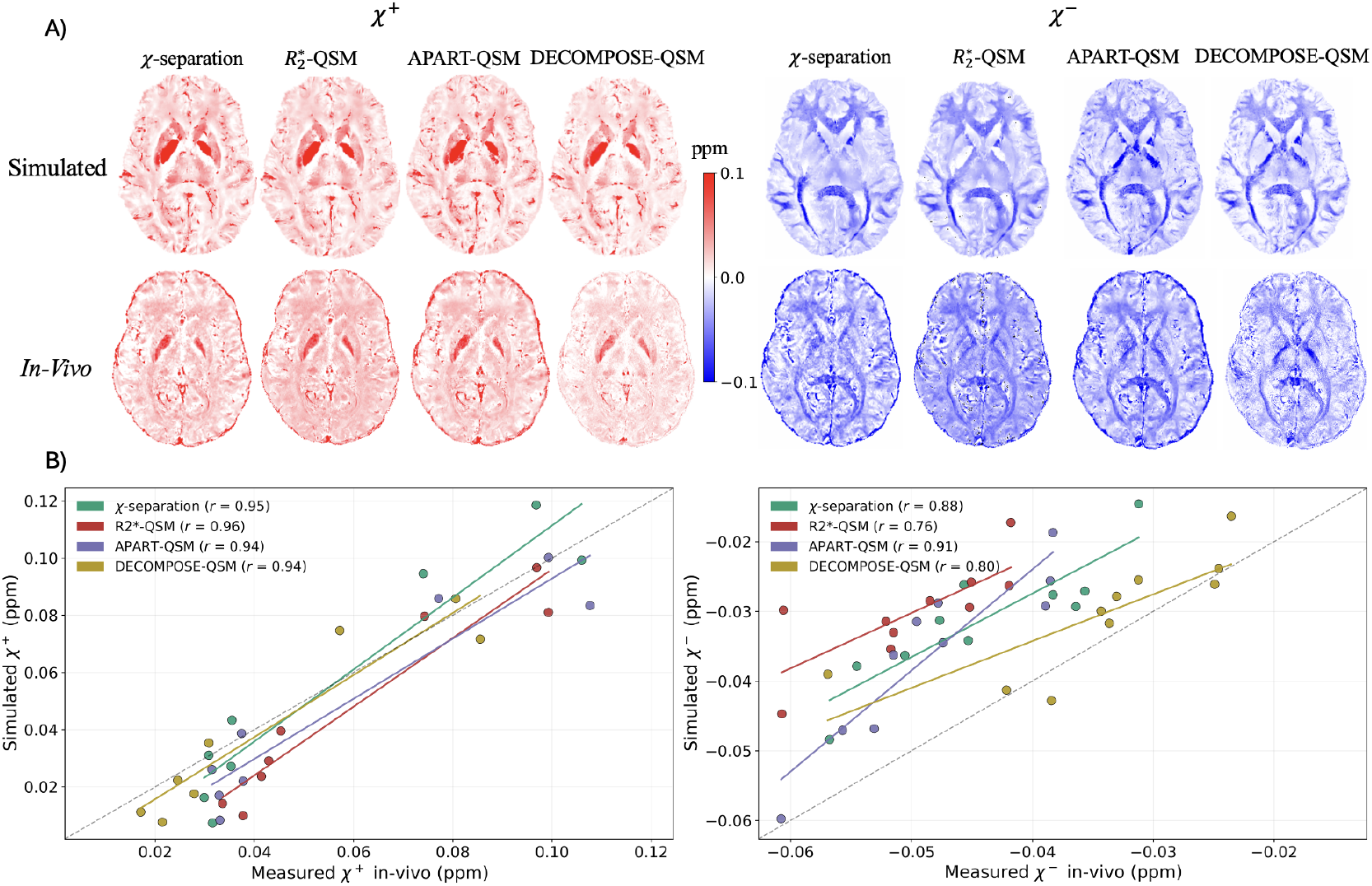
(A) Simulated and in-vivo measured susceptibility maps for χ^+^ (left) and χ^−^ (right) components across susceptibility separation methods: χ-separation, 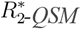, APART-QSM, and DECOMPOSE-QSM. The color scale indicates susceptibility values in ppm, with red representing positive susceptibility and blue representing negative susceptibility. (B) Simulated vs. in-vivo measured susceptibility values as a scatter plot for χ^+^ (left) and χ^−^ (right) across the four susceptibility separation methods. Susceptibility values were averaged across predefined ROIs. The dashed line represents y = x as a reference and the correlation coefficients are in the plot’s legend.

The simulated and in-vivo local field maps also displayed visually similar patterns (Figure 7). Quantitatively, the scatter plot in the right panel of Figure 7 shows a high correlation between the simulated and in-vivo measured values averaged across WM sub-ROIs (Table 1, column 2), with a correlation coefficient of 0.91 and a slope of 1.

**Figure 7.**
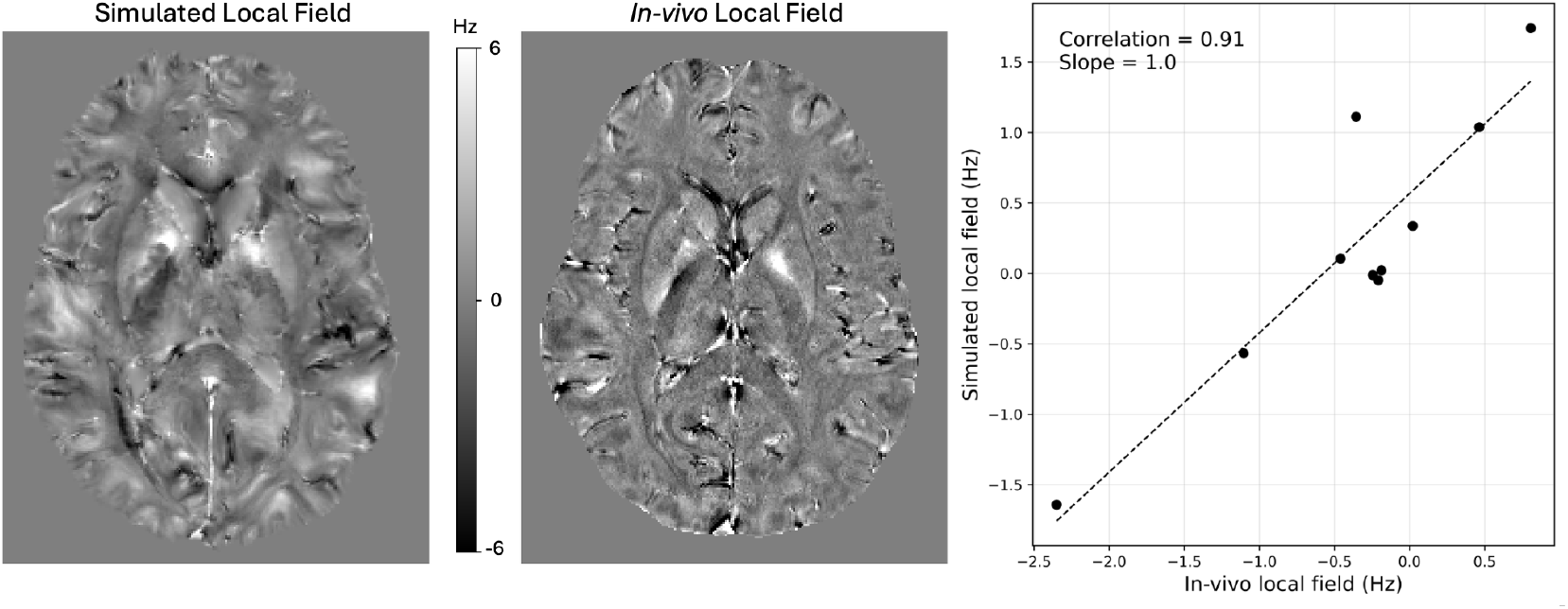
Comparison of simulated and in-vivo local field maps. The left panel displays the simulated local field map, while the middle panel shows the in-vivo local field map. The right panel presents a scatter plot (each datapoint represents a WM sub-ROI) of simulated versus in-vivo local field values, with a dashed linear regression line.

Finally, Figure 8 shows the comparison between the generated *T*_2_ map and the *T*_2_ map measured from in-vivo data. Visually, the two maps exhibit high similarity, with comparable contrast across different brain regions. Simulated vs. in-vivo measured *T*_2_ values averaged within various ROIs revealed a strong correlation (*r*=0.96) and a slope of 0.95.

**Figure 8.**
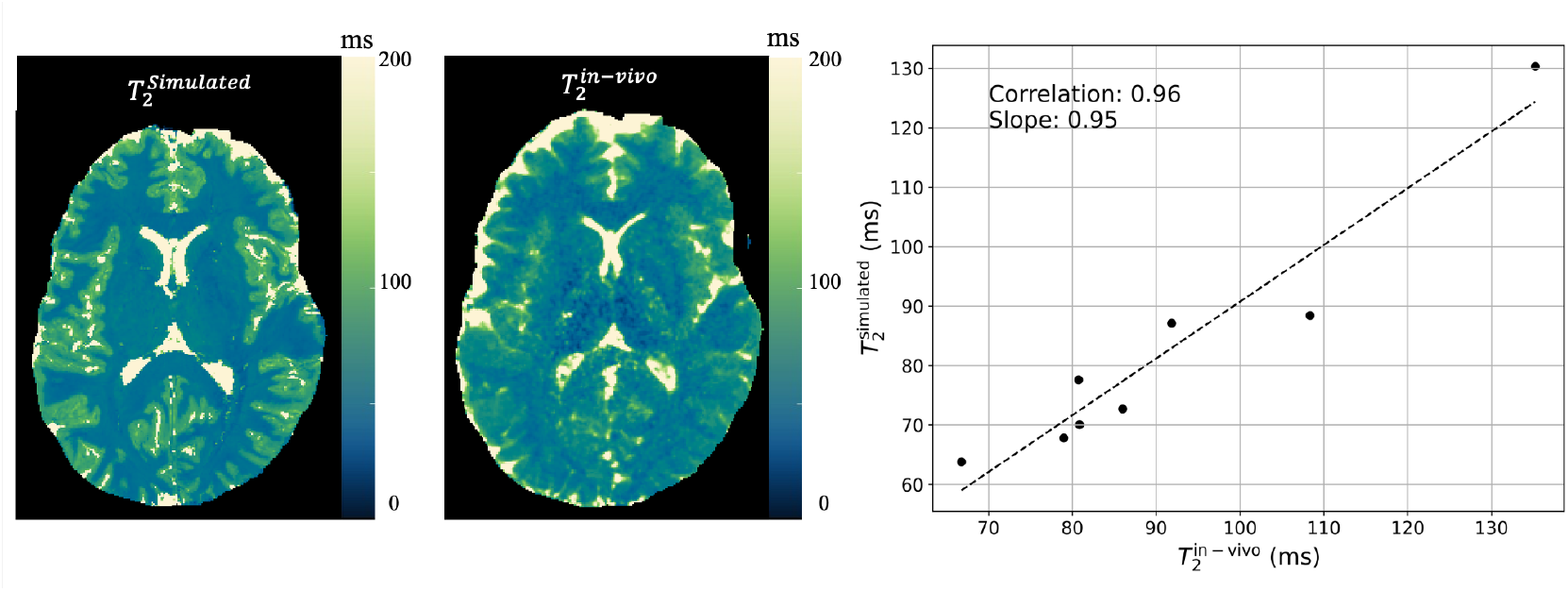
The left panel: Generated 3 T T_2_ map. Middle panel: In-vivo 3 T T_2_ map. Right panel: Scatter plot of the simulated vs. in-vivo T_2_ values extracted from predefined ROIs . The dashed line represents the linear regression line.

## Discussion

The original QSM validation phantom was developed to address the growing number of QSM methods, providing a way to directly test and validate these algorithms in a controlled environment with known numerical ground truth values. Given the growing interest in susceptibility separation techniques and the emergence of new algorithms, a dedicated phantom provides a valuable framework for systematically assessing these methods. In this study, we developed an in-silico brain phantom, inspired by the QSM phantom, that incorporates both positive (paramagnetic) isotropic and negative (diamagnetic) isotropic or anisotropic susceptibility sources. This phantom can be used by the research community to simulate gradient echo data under defined conditions and to examine how susceptibility separation algorithms respond to controlled changes in the phantom, using its known numerical ground truth as a reference.

In this study, we used our newly developed phantom to investigate the impact of susceptibility anisotropy on four susceptibility separation algorithms: χ-separation, 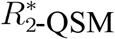, APART-QSM, and DECOMPOSE-QSM. These algorithms do not account for susceptibility anisotropy effects in white matter. We aimed to quantify errors associated with neglecting anisotropic contributions to susceptibility by comparing results obtained with phantom versions that do vs. do not include susceptibility anisotropy. Our findings showed that, depending on the algorithm, the MSPE in χ^−^ estimates increased by up to 53% when anisotropy effects were present. Moreover, the error associated with χ^−^ varied by up to ∼57% when WM fibers were parallel vs. perpendicular to *B*_0_. Together, these results indicate that susceptibility anisotropy can meaningfully influence susceptibility separation results and should not be neglected.

The proposed phantom is built upon a simplified representation of tissue susceptibility sources and signal behavior. This includes the adoption of the static dephasing regime, which is often reasonable in WM at TE ≲ 40 ms^41,49,50^, but it can break down near the microvasculature where diffusion-narrowing effects modulate MGRE/MESE signals, violating static-dephasing^51^. More generally, since the phantom does not necessarily reproduce each biophysical feature assumed by every susceptibility separation model, disagreement between an algorithm’s estimates and the phantom ground truth should not always be interpreted as evidence of reduced algorithmic accuracy; in some cases, it may instead reflect a mismatch between the assumptions used to construct the phantom and those underlying the algorithm being evaluated. While our digital phantom and simulation framework provide a controlled numerical environment for examining algorithm behavior under defined conditions, including the presence vs. absence of susceptibility anisotropy and across different SNR levels; results should be interpreted in light of the assumptions made in both our phantom and simulations framework and the fitting model in question. This issue is less pronounced in the original QSM in-silico phantom because QSM methods typically share the same dipole-based forward model and differ mainly in inversion strategy and regularization, whereas susceptibility separation methods can differ more fundamentally in their signal-model assumptions.

The limitation associated with the fitting vs. phantom model assumptions is illustrated by DECOMPOSE-QSM, which estimates paramagnetic, diamagnetic, and neutral susceptibility components, whereas the present phantom includes only paramagnetic and diamagnetic sources. The higher χ^−^ MSPE observed for DECOMPOSE-QSM may therefore reflect, at least in part, a mismatch between the algorithm’s signal model and the phantom design, rather than inferior performance. More broadly, susceptibility anisotropy did not affect all algorithms to the same extent. 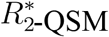 and DECOMPOSE-QSM, both of which rely solely on 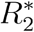, showed the smallest increase in χ^−^ MSPE (10% and 13%, respectively) when anisotropy was introduced. However, both of those algorithms also had the highest MSPE in the no anisotropy condition. At SNR = 300, DECOMPOSE-QSM also exhibited the smallest variation in the MSPE of parallel vs. perpendicular fibers. These findings indicate that algorithms that do not measure 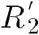 data are less sensitive to myelin’s magnetic behavior. APART-QSM was the algorithm that resulted in the lowest mean MSPE for simulations with anisotropy effects. However, this improved performance may be partly attributed to the iterative approach used by APART-QSM for computing D_r_, which may implicitly account for some anisotropic effects. χ-separation, 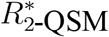, and DECOMPOSE-QSM, on the other hand, used a constant D_r_ value across the entire brain. When considering the MEV at SNR = 300 for MSPE of parallel vs. perpendicular fibers, χ-separation had the highest value (i.e., greatest sensitivity to anisotropy effects). The impact of GRE noise on the sensitivity to susceptibility anisotropy was also examined. Higher GRE noise levels reduced the ability to discern the influence of orientation angle on the measured susceptibility. At SNR = 50, the MEV for APART-QSM was the lowest at 9.9%.

We also examined how varying levels of GRE SNR affect the performance of the four susceptibility separation methods when it comes to estimating both χ^+^ and χ^−^. Our results showed that at high SNR, APART-QSM had the lowest MSPE for χ^−^and χ-separation had the lowest MSPE for χ^+^. However, as SNR increased from 50 to 300, APART-QSM showed the greatest change in MSPE (an 84% and 82% decrease for χ^+^ and χ^−^, respectively), and DECOMPOSE-QSM showed the second greatest change (a 72% and 57% decrease for χ^+^ and χ^−^, respectively). The pronounced sensitivity of APART-QSM to SNR may be attributed to its iterative algorithm for calculating D_r_^52^. Furthermore, APART-QSM and DECOMPOSE-QSM lack explicit weighting factors for noise stabilization, unlike χ-separation, which incorporates a weighting factor that accounts for the GRE signals’ SNR. χ-separation was the most stable across SNR, with a 58% and 10% decrease in MSPE for χ^+^ and χ^−^, respectively, and the lowest MSPE when averaged across all SNR levels, for both χ^+^ and χ^−^.

While χ^+^ values at high SNR had relatively low MSPEs (∼10-20%), further investigations revealed that this was driven by low MSPEs in GM regions (see supplementary materials, Figure S3). χ^+^ MSPE values in WM regions were very high (Figure S2), ranging from ∼3600% to ∼6200%. This underscores the difficulty in accurately estimating χ^+^ in WM, where the values of susceptibility are small (on the order of 1 to 10 ppb), and the potential limited value of χ^+^-based analyses in WM. Notably, across all algorithms, no statistically significant differences (*p* > 0.05) were observed between the with and without susceptibility anisotropy conditions, indicating that large errors persist in WM regardless of anisotropy.

The phantom was designed to mimic in-vivo conditions to ensure the realism of the simulated data. The phantom includes a realistic *T*_2_ map, which is required for some susceptibility separation techniques. While *T*_2_ remains relatively stable across clinical field strengths, it tends to decrease at ultra-high *B*_0_^38,53^. Given the challenges of establishing a precise model for *T*_2_’s dependence on *B*_0_^38,53^ and the prevalence of *T*_2_ measurements at 3 T, we decided to take a scaling approach to create a 7 T *T*_2_ map from a generated 3 T *T*_2_ map. We verified our generated 3 T *T*_2_ map by comparing it with a 3 T *T*_2_ map obtained in-vivo and found excellent agreement between the two. However, the 7 T generated *T*_2_ map was not verified using in-vivo measurements due to the challenges associated with 7 T *T*_2_ mapping, such as high specific absorption rate (SAR) and *B*_1_ field inhomogeneities^54^. The phantom also includes a 3 T *T*_1_ map that was created by scaling the original QSM validation phantom’s 7 T *T*_1_ map to 3 T. Although we didn’t acquire an in-vivo 3 T *T*_1_ map, the generated 3 T *T*_1_ map was verified using reported *T*_1_ values from Gelman *et al*.^55^. We found a strong correlation (*r* = 0.99) and a slope of 0.94 between the generated 3 T *T*_1_ values and those literature values. To verify the phantom’s χ^+^ and χ^−^ maps, MGRE and MESE data were acquired in a healthy volunteer and used to obtain measured χ^+^ and χ^−^ maps. We found a strong agreement between the in-vivo measured χ^+^ and χ^−^ (for all four algorithms) and the χ^+^ and χ^−^ values measured from data simulated with our phantom (correlation values for χ^−^ ranged from 0.94 to 0.96 and from 0.76 to 0.91 for χ^−^). Although the simulated and in-vivo measurements showed strong agreement, it should be noted that the phantom adopts a single-compartment relaxation model, and therefore cannot model compartment-specific *T*_1_-weighting. However, the relatively short TR used in our in-vivo MGRE acquisition may have led to incomplete longitudinal recovery and, therefore, *T*_1_-dependent compartment weighting, an effect that is not represented in the phantom. Consequently, part of any mismatch between simulated and in-vivo measurements may be attributed to these acquisition-dependent features.

To further assess the anisotropy model used in our study, we compared a measured in-vivo local field map to one generated from data simulated using our phantom. Despite the strong correlation between in-vivo and simulated local field values, the contrast in our simulated maps differed in some regions from that observed in-vivo. This discrepancy may be due to the fact that our phantom does not account for microstructural anisotropy. Work carried out at 7 T by Wharton and Bowtell^56^ showed that when gradually introducing different sources of anisotropy into local field simulations, incorporating only susceptibility anisotropy produced relatively low contrast between regions, but adding microstructural anisotropy yielded simulated local field maps that more closely matched in-vivo measurements. They also showed that the induced frequency offsets due to anisotropic susceptibility were much smaller in magnitude than the frequency contribution due to microstructure. The existing QSM validation phantom has the option of simulating frequency offsets caused by microstructure anisotropy by using a theoretical model^57^ with coefficients obtained from 7 T measurements carried out by Wharton and Bowtell^56^. Here, we sought to focus on evaluating the effects of susceptibility anisotropy on susceptibility separation techniques. However, Wharton and Bowtell’s^56^ results suggest that the additional inclusion of microstructure effects in our phantom may further impact susceptibility separation results, which warrants further study. The microstructure anisotropy model included in the QSM validation phantom can also be used for 7 T simulations with our phantom. Extending its use to 3 T could be achieved by scaling the model’s coefficients to 3 T. While scaling the model’s coefficients provides a straightforward initial approach, repeating the experiments proposed by Wharton and Bowtell’s at 3 T would result in more accurate coefficients.

## Conclusion

The possibility of accurately quantifying positive and negative magnetic susceptibility has garnered considerable interest within the community. Within the last five years, several models^13–16^ have been proposed. However, none of these explicitly account for myelin’s magnetic susceptibility anisotropy. In this work, we have proposed an open-source brain tissue phantom that enables controlled comparison of different susceptibility separation scenarios against a known numerical ground truth within a defined simulation framework. This is currently lacking in the field, making it difficult to compare between susceptibility separation models. To demonstrate our phantom’s benefits, we have used it to study the impact of magnetic susceptibility anisotropy on susceptibility separation results. Our results suggest that anisotropy has a non-negligible impact on susceptibility separation results. These findings underscore the importance of developing new algorithms that incorporate anisotropic effects to achieve more accurate and reliable susceptibility mapping.

## Data and code availability

The code used to create susceptibility and relaxation maps, and simulate GRE signals is available at: https://github.com/neuropoly/Susceptibility-Separation-Phantom, while the data used to generate the figures described in this paper can be found here: https://osf.io/9xwhz/.

## Acknowledgments

This work is supported by the TransMedTech Institute, thanks to the financial support of the Canada First Research Excellence Fund and the Fonds de recherche du Québec, the Natural Sciences and Engineering Research Council of Canada (NSERC), and Polytechnique Montreal.

## Supplementary Material

Supplementary material is available.

