## Supplementary Material for "A realistic in-silico brain phantom for quantifying susceptibility anisotropy-induced error in susceptibility separation"

1) **Values of** $C$ and $\beta$

The $T_1$ model parameters (C and β) were determined using a least squares fitting approach based on the values reported by Rooney et al. [38] for multiple brain regions. The results, presented in Table 1, show that $T_1$ values generally increase with B0, consistent with known tissue relaxation behavior. However, the CSF appears to be an exception, with its $T_1$ remaining approximately constant across field strengths, as indicated by a near-zero $\beta$ value.

| **ROIs** | **$T_1$ model** |
|---|---|
| Caudate nucleus | $T_1=0.954(B_0)^{0.325}$ |
| Globus pallidus | $T_1=0.664(B_0)^{0.367}$ |
| Putamen | $T_1=0.855(B_0)^{0.352}$ |
| Red nucleus | $T_1=0.459(B_0)^{0.508}$ |
| Dentate nucleus | $T_1=0.459(B_0)^{0.508}$ |
| Substantia nigra | $T_1=0.459(B_0)^{0.508}$ |
| Thalamus | $T_1=0.817(B_0)^{0.357}$ |
| White matter | $T_1=0.583(B_0)^{0.376}$ |
| Grey matter | $T_1=0.857(B_0)^{0.376}$ |
| CSF | $T_1=4.322(B_0)^{-0.006}$ |

*Table S1: $T_1$ model for QSM validation phantom ROIs obtained by a least square fit of the model proposed by Rooney et al., [38].*

### 2) 3 T vs 7 T Bland Altman

Figure S2 shows Bland-Altman analyses of the measured 3 T and 7 T $\chi^{-}$ in WM sub-ROIs with susceptibility anisotropy effects. The plots indicate a bias of 0.0034, -0.0024, 0.0057, and -0.0109 ppm for $\chi$-separation, $R_2^*$-QSM, APART-QSM, and DECOMPOSE-QSM, respectively. Overall, the Bland-Altman analysis shows a weak systematic difference as most points are within the lines of agreement, with a more pronounced bias in DECOMPOSE-QSM.

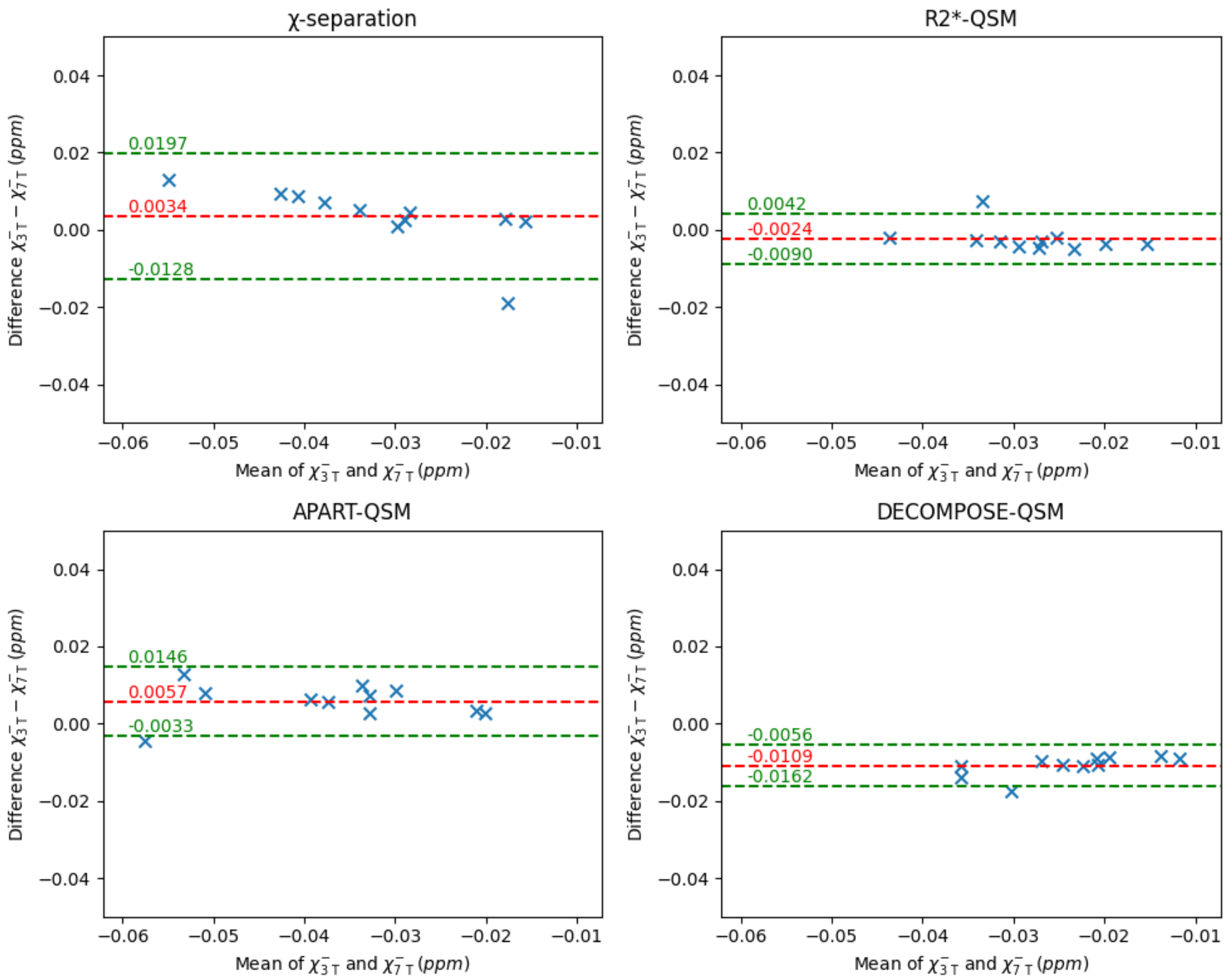


*Figure S1: Bland–Altman plot comparing the measured $\chi^{-}$ values using four susceptibility separation techniques from 3 T and 7 T simulated data with susceptibility. Each data point represents the averaged susceptibility value within a WM sub-ROI.*

#### 3) $\chi^+$ simulated vs measured

Figure S2 shows kernel density estimate (KDE) plots of the WM sub-ROI-averaged MSPE values for $\chi^+$ measured from noiseless 3 T data simulated with and without susceptibility anisotropy effects. For algorithms that use $R_2'$ ($\chi$-separation and APART-QSM), the MSPE is higher when susceptibility anisotropy is included. However, for algorithms that use $R_2^*$ (DECOMPOSE-QSM and $R_2^*$-QSM), the MSPE is lower when susceptibility anisotropy is included. MSPE values for $\chi^+$ are considerably large, highlighting the limitations of these methods to the small susceptibility values encountered in WM in $\chi^+$.

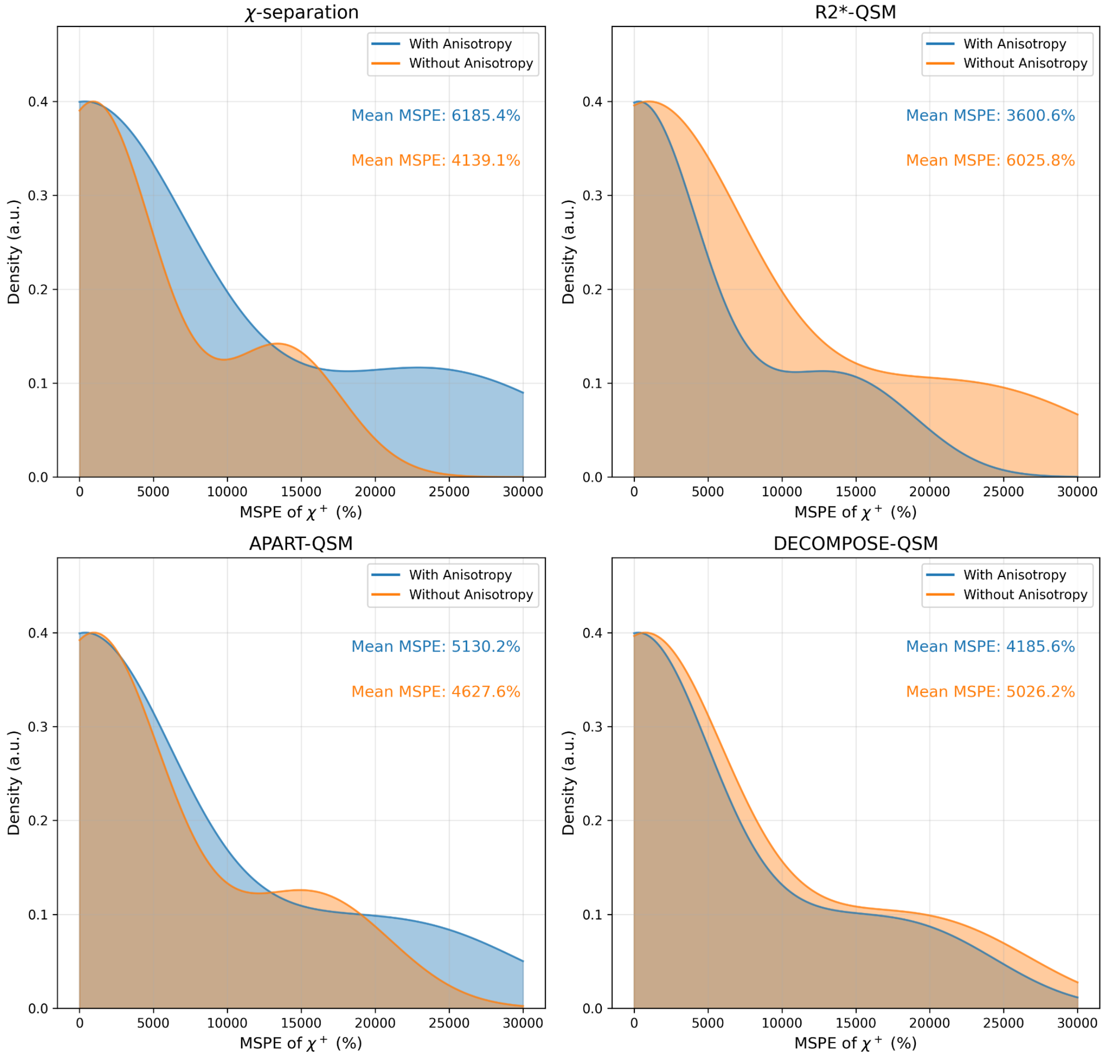


*Figure S2: KDE plots of the WM sub-ROI-averaged MSPE of $\chi^+$ measured from noiseless 3 T data simulated with (blue) and without (orange) susceptibility anisotropy using four susceptibility separation techniques: $\chi$-separation, $R_2^*$-QSM, APART-QSM, and DECOMPOSE-QSM.*

Figure S3 shows KDE plots of the QSM validation ROI-averaged (without WM) MSPE values for $\chi^+$ measured from noiseless 3 T data simulated with and without susceptibility anisotropy effects. $\chi$-separation achieved the lowest MSPE value, followed by APART-QSM, then $R_2^*$-QSM, and finally DECOMPOSE-QSM. For every algorithm tested, the distribution of MSPE with anisotropy appears to be broader and has a higher average than that of the case without anisotropy indicating that although susceptibility anisotropy was not modeled in GM it still affects the surrounding tissue.

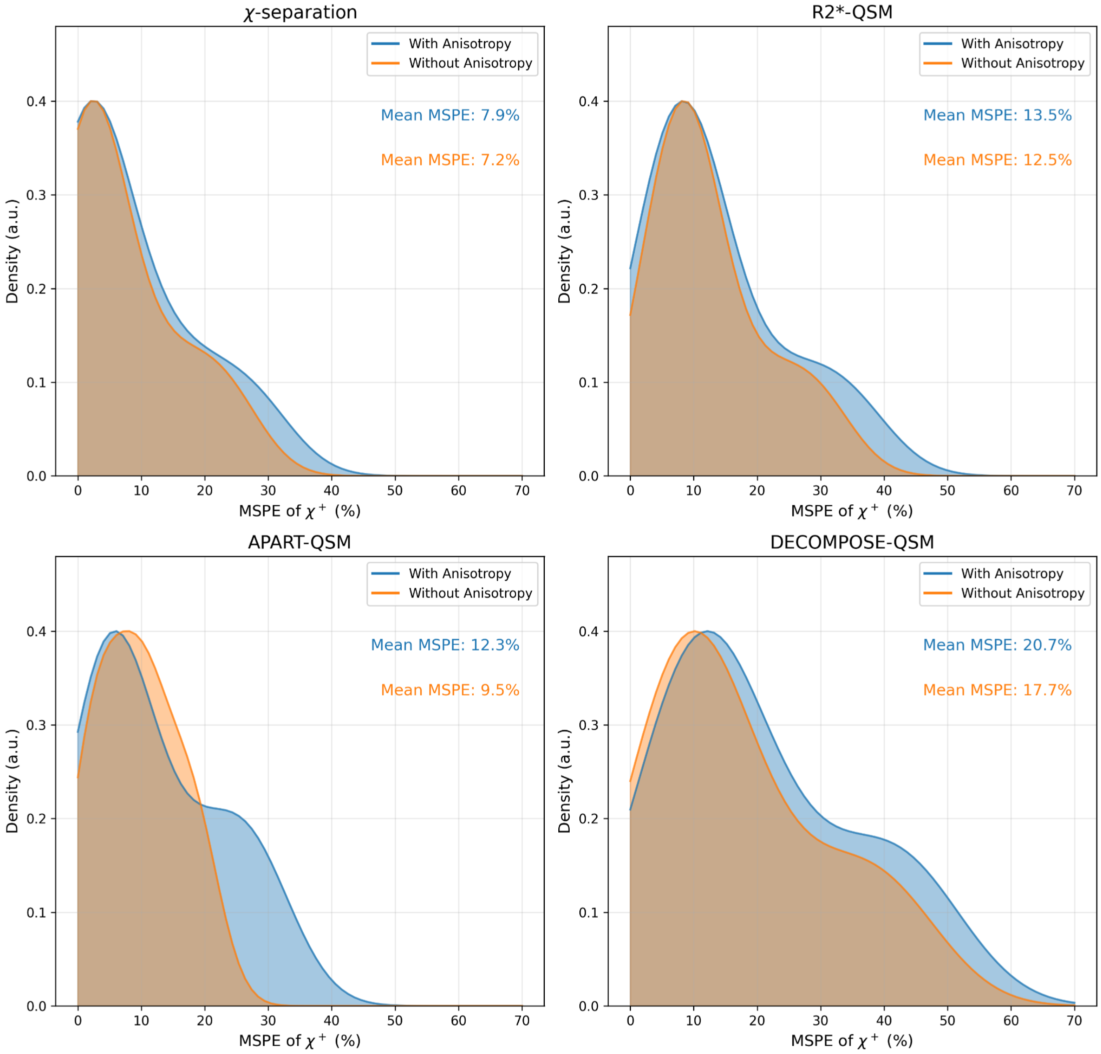


*Figure S3: KDE plots of the QSM validation ROI-averaged MSPE of $\chi^+$ measured from noiseless 3 T data simulated with (blue) and without (orange) susceptibility anisotropy using four susceptibility separation techniques: $\chi$-separation, $R_2^*$-QSM, APART-QSM, and DECOMPOSE-QSM.*
